# Checkpoint defects require WRNIP1 to counteract R-loop-associated genomic instability

**DOI:** 10.1101/858761

**Authors:** Veronica Marabitti, Giorgia Lillo, Eva Malacaria, Valentina Palermo, Pietro Pichierri, Annapaola Franchitto

## Abstract

Conflicts between replication and transcription are common source of genome instability and many factors participate in prevention or removal of harmful R-loops. Here, we demonstrate that a WRNIP1-mediated response plays an important role in counteracting accumulation of aberrant R-loops. Using human cellular models with compromised ATR-dependent checkpoint activation, we show that WRNIP1 is stabilised in chromatin and is needed for maintaining genome integrity by mediating the ATM-dependent phosphorylation of CHK1. Furthermore, we show that loss of WRN or ATR signalling leads to accumulation of R-loop-dependent parental ssDNA, which is covered by RAD51. We demonstrate that WRNIP1 chromatin retention is also required to stabilise the association of RAD51 with ssDNA in proximity of R-loops. Therefore, in these pathological contexts, ATM inhibition or WRNIP1 abrogation is accompanied by increased levels of genomic instability. Overall our findings reveal a novel function of WRNIP1 in preventing R-loop-driven genome instability, providing new clues to understand the way replication-transcription conflicts are resolved.

## INTRODUCTION

DNA damage or unusual DNA structures may pose a serious impediment to DNA replication. One of the major obstacles to the replication fork progression is transcription (Aguilera & García-Muse, 2013; Gorgoulis *et al*, 2005). Encounters between replication and transcription machineries are unavoidable, as they compete for the same DNA template, so collisions occur frequently (García-Muse & Aguilera, 2016; Bermejo *et al*, 2012). The main transcription-associated structures that can be detrimental to fork movement are R-loops (Allison & Wang, 2019; Gaillard & Aguilera, 2016). They are transient and reversible structures forming along the genome, consisting of a DNA-RNA hybrid and a displaced single-stranded DNA. Despite their beneficial function in a series of physiological processes, such as transcription termination, regulation of gene expression and DNA repair ((Aguilera & García-Muse, 2012), if their turnover is deregulated, they can cause a head-on clash between the replisome and the RNA polymerase leading to R-loop-driven replication stress (Gómez-González & Aguilera, 2019; Crossley *et al*, 2019). Therefore, R-loop accumulation is a leading source of replication stress and thus genome instability. As critical levels of R-loops may contribute to the heightened cancer predisposition of humans, cells have evolved multiple factors to prevent/remove these harmful structures. Apart from the canonical mechanisms, a lot of evidences have highlighted the importance of several replication fork protection factors and DNA damage response (DDR) proteins in counteracting pathological R-loop persistence. Among them are BRCA1 and 2 (Bhatia *et al*, 2014; Shivji *et al*, 2018; Hatchi *et al*, 2015), the components of the Fanconi Anaemia pathway (Schwab *et al*, 2015; García-Rubio *et al*, 2015; Liang *et al*, 2019), DNA helicases RECQ5 and BLM (Urban *et al*, 2016; Chang *et al*, 2017) and the apical activator of the DDR, the ATM kinase (Tresini *et al*, 2015). Interestingly, defects in the ATR-CHK1 signalling promote accumulation of aberrant R-loops (Barroso *et al*, 2019). Recently, a novel function in the response to R-loop-associated genome instability in human cells has been reported for the Werner syndrome protein (WRN) (Marabitti *et al*, 2019).

WRN is a protein belonging to the RecQ family of DNA helicases essential in genome stability maintenance, with a role in the repair and recovery of stalled replication forks (Pichierri *et al*, 2011; Rossi *et al*, 2010). WRN-deficient cells show impaired ATR-dependent checkpoint activation after short times of treatment with aphidicolin-induced mild replication stress (Basile *et al*, 2014). Among the plethora of WRN-interacting proteins that participate in the maintenance of genome stability, there is the still poorly characterized human Werner helicase interacting protein 1 (WRNIP1) (Kawabe Yi *et al*, 2001; Kawabe *et al*, 2006). WRNIP1 is a member of the AAA+ class of ATPase family that is evolutionary conserved (Kawabe Yi *et al*, 2001; Hishida *et al*, 2001). Although the yeast homolog of WRNIP1, Mgs1, is required to prevent genome instability caused by replication arrest (Branzei *et al*, 2002), little is known about the function of human WRNIP1. Previous studies established that WRNIP1 binds to forked DNA that resembles stalled forks (Yoshimura *et al*, 2009). More recently, we demonstrated that WRNIP1 is recruited to hydroxyurea-induced stalled replication forks where it interacts with RAD51 (Leuzzi *et al*, 2016). Indeed, WRNIP1 is directly involved in preventing uncontrolled MRE11-mediated degradation of stalled forks by promoting RAD51 stabilization on single-stranded DNA (ssDNA) (Leuzzi *et al*, 2016). Furthermore, WRNIP1 has been implicated in the activation of the ATM-dependent checkpoint in response to mild replication stress (Kanu *et al*, 2016).

In this study, we investigated a possible role of the WRN-interacting protein 1 (WRNIP1) in the response to aberrant R-loop accumulation in cells with compromised replication checkpoint response, and how it prevents excessive levels of DNA damage and the consequent genomic instability upon mild replication stress. We find that loss of WRN or depletion of essential factors for ATR-CHK1 pathway activation promotes WRNIP1 retention in chromatin. Furthermore, we provide evidence that, in these pathological contexts, WRNIP1 plays a dual role: to maintain genome integrity mediating the ATM-dependent phosphorylation of CHK1 as well as stabilising the association of RAD51 with ssDNA in proximity of R-loops after mild replication stress.

## RESULTS

### Combined loss of WRNIP1 and WRN results in increased sensitivity of cells to mild replication stress

WRNIP1 was originally identified as a WRN-interacting protein (Kawabe Yi *et al*, 2001), but there is no evidence that they cooperate in response to mild replication stress. Hence, we first investigated if WRN and WRNIP1 interact in vivo testing their co-immunoprecipitation. To this aim, HEK293T cells were transfected with the FLAG-tagged wild-type WRNIP1 and treated or not with low-dose of aphidicolin (Aph). Under untreated conditions, WRNIP1 and WRN co-immunoprecipitated, as expected, and Aph slightly increased this interaction (Figure 1A). This result supports a possible cooperation of WRNIP1 and WRN in response to mild replication stress.

**Figure 1.**
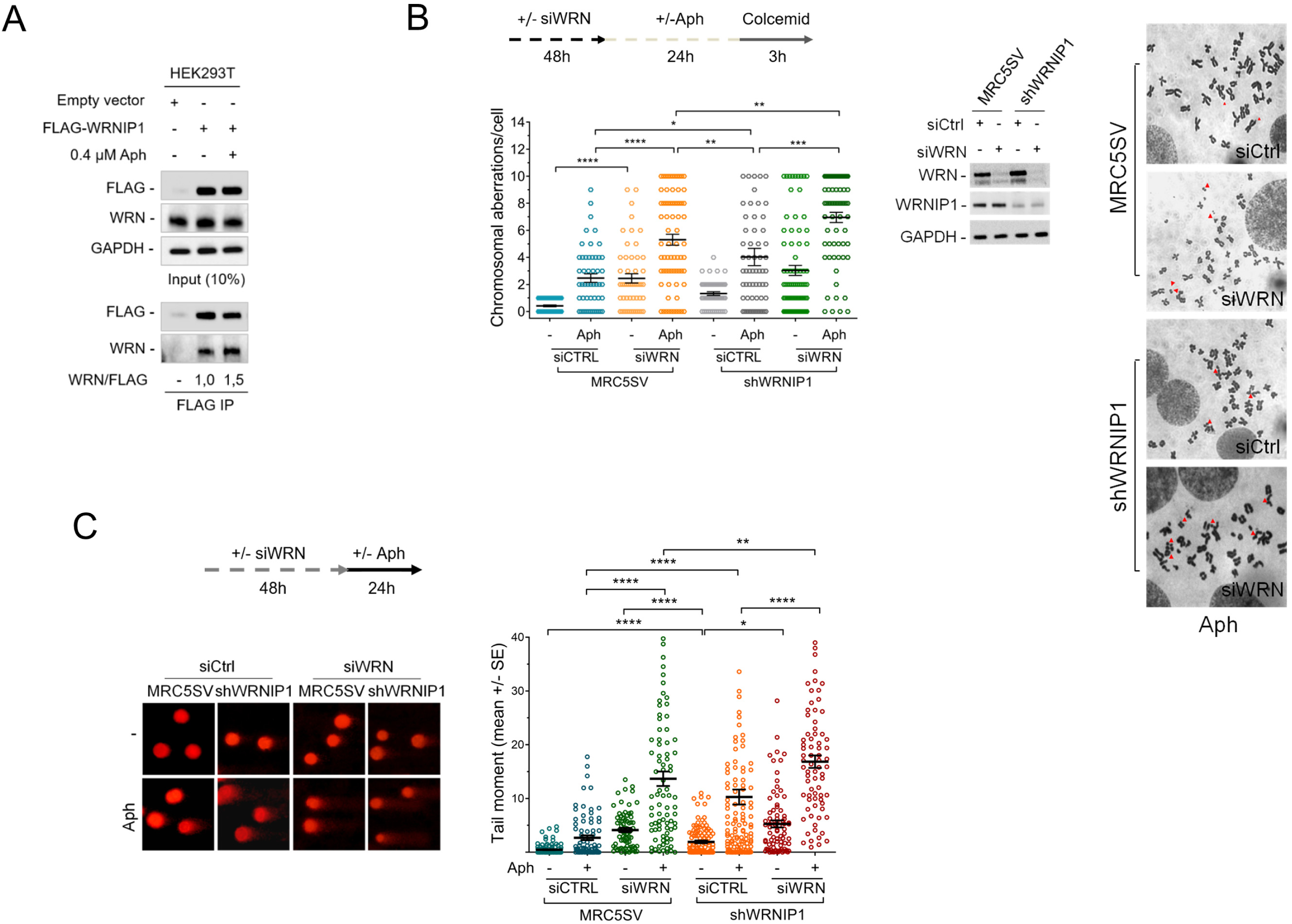
Enhanced genomic instability in cells lacking both WRNIP1 and WRN. **A)** Co-immunoprecipitation experiments in HEK293T cells transfected with empty vector or FLAG-WRNIP1 plasmid. Cells were treated or not with 0.4 μM Aphidicolin (Aph) for 24 hours. After treatment, cell lysates were immunoprecipitated (FLAG IP) using anti-FLAG antibody. The presence of WRNIP1 and WRN was assessed by immunoblotting using the anti-FLAG and anti-WRN antibody, respectively. Whole cell extracts were analysed (Input 10%). The membrane was probed with the same antibodies used for IP. GAPDH was used as a loading control. **B)** Analysis of chromosomal aberrations in wild-type fibroblasts (MRC5SV) or MRC5SV cells stably expressing shRNA against WRNIP1 (shWRNIP1) transfected with siRNAs directed against GFP (MRC5SV or shWRNIP1) or WRN (MRC5SV ^siWRN^ or shWRNIP1^siWRN^). Experimental scheme is reported. Scatter dot plot shows data presented as chromosomal aberration per cell. Error bars represent standard error (* P < 0.1; ** P < 0.01; *** P < 0.001; **** P < 0.0001; two-tailed Student’s t test). Representative Giemsa-stained metaphases are given. Arrows indicate chromosomal aberrations. Down-regulation of WRN and WRNIP1 was verified by immunoblotting using the relevant antibodies. GAPDH was used as a loading control. **C)** Analysis of DNA breakage accumulation evaluated by alkaline Comet assay. Experimental scheme is reported. Cells were transfected as in (B) before treatment with Aph and then subjected to alkaline Comet assay. Scatter dot plot shows data presented as tail moment +/- SE. Error bars represent standard error (** P < 0.01; **** P < 0.0001; two-tailed Student’s t test). Representative images are given.

WRN-deficient cells are hyper-sensitive to Aph (Pirzio *et al*, 2008). Hence, to gain insight into the role of WRNIP1 and WRN in maintaining genome stability upon mild replication stress, we examined the level of chromosomal damage in cells lacking both proteins. MRC5SV cells stably expressing WRNIP1-targeting shRNA (shWRNIP1) (Leuzzi *et al*, 2016) and their parental, wild-type, counterpart (MRC5SV) were transiently transfected with WRN-targeting siRNA. After transfection, cells were exposed to Aph and aberrations scored in metaphase chromosomes. As previously reported (Pirzio *et al*, 2008), depletion of WRN determined an increase in the average number of gaps and breaks both in unperturbed and Aph-treated samples, respect to wild-type cells (Figure 1B). On the contrary, shWRNIP1 cells presented a level of chromosomal damage that is slightly higher than in wild-type cells, but lower respect to WRN-depleted cells (Figure 1B). Notably, concomitant depletion of WRNIP1 and WRN resulted associated with a synergistic enhancement of the chromosomal aberration frequency (Figure 1B).

In parallel experiments, we also evaluated the presence of DNA damage by alkaline Comet assay. Loss of WRNIP1 or WRN led to higher spontaneous levels of DNA damage compared to wild-type cells, and depletion of WRN in shWRNIP1 cells significantly increased DNA damage accumulation respect to each single deficiency (Figure 1C). Similarly, concomitant depletion of WRN and WRNIP1 increased DNA damage above the levels observed in single-depleted cells after Aph treatment (Figure 1C).

Overall, these findings indicate that WRNIP1-deficiency sensitises cells to Aph and that combined loss of WRNIP1 and WRN exacerbates this sensitivity. Furthermore, they suggest that these two proteins do not function cooperatively in response to mild replication stress, even if their association in a complex is stimulated by Aph.

### WRNIP1 is tightly associated with chromatin in WS cells

Our results argue that WRNIP1 and WRN cooperate to maintain genome integrity upon mild replication stress, although they do not act in the same pathway. Hence, we verified whether WRNIP1 might be required in the absence of WRN. Since protein recruitment/retention onto chromatin is a critical process for DNA metabolism, we monitored the chromatin association of WRNIP1 in WRN-deficient cells (WS) and in isogenic WRN wild-type corrected (WSWRN) counterpart. Cells were treated or not with Aph and subjected to a chromatin fractionation assay at increasing concentrations of NaCl combined with detergent pre-extraction (Figure 2A). WRNIP1 total levels were comparable in WS and WS-corrected cell lines under unperturbed and Aph-treatment conditions (Figure 2B). Although low-salt extraction did not greatly influence WRNIP1 binding to chromatin, under high-salt concentration, chromatin-bound fraction of the protein was higher in WS than in WS-corrected cells (Figure 2B). Similar results were obtained in cells transiently-depleted of WRN (WRN-kd), indicating that increased stability of WRNIP1 in chromatin is not cell type-dependent but related to the absence of WRN (Suppl. Figure 1). Our findings suggest that loss of WRN increases the affinity of WRNIP1 for chromatin.

**Figure 2.**
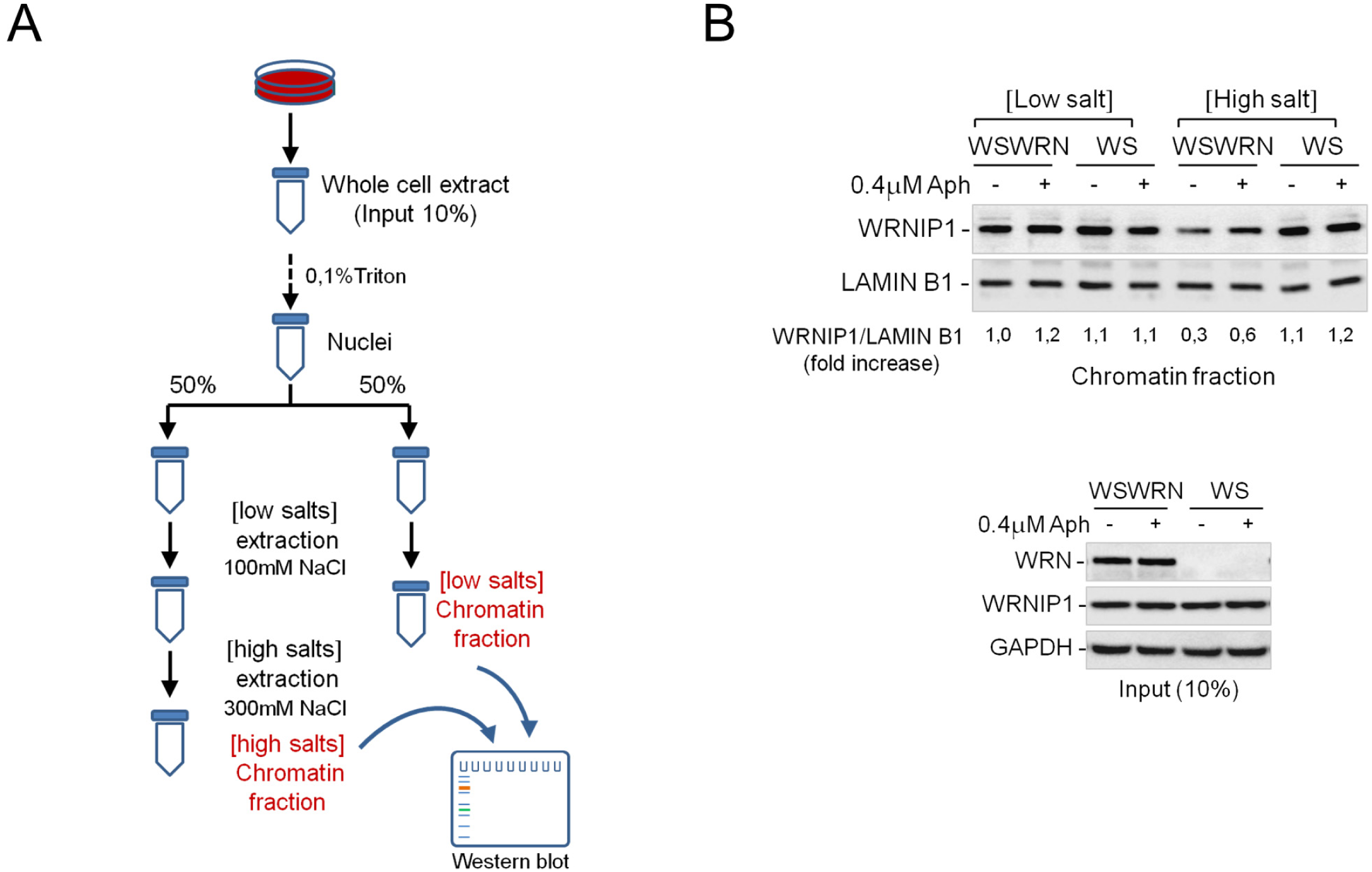
WRN absence increases WRNIP1 retention onto chromatin. **(A)** Schematic representation of chromatin fractionation analysis procedure (see also “Materials and Methods” section). **B)** WB analysis of chromatin binding of WRNIP1 in WSWRN and WS cells treated or not with Aph. Chromatin fractionation was performed under low or high salt concentrations. The membrane was probed with an anti-WRNIP1 antibody. LAMIN B1 was used as a loading for the chromatin fraction. Total amount of WRNIP1 (Input 10%) in the cells was determined with the relevant antibody. GAPDH was used as a loading control. The amount the chromatin-bound WRNIP1 is reported as ratio of WRNIP1/LAMIN B1 normalized over the untreated control.

### WRNIP1 mediates ATM signalling activation leading to CHK1 phosphorylation in response to mild replication stress in WS cells

WRNIP1 promotes ATM signalling activation upon mild replication stress (Kanu *et al*, 2016), and we recently reported a hyper-activation of ATM in WRN-deficient cells after mild-replication stress (Marabitti *et al*, 2019). To elucidate the significance of the higher chromatin-affinity of WRNIP1 in WS cells, we evaluated whether it correlates to ATM activation. Hence, we analysed by immunofluorescence the presence of the phosphorylation of ATM at Ser1981 (pATM), an accepted marker for ATM activation (Bakkenist & Kastan, 2003), in wild-type (WSWRN) or WS cells depleted of WRNIP1 treated or not with Aph. As expected, pATM levels were increased in the absence of WRN (Figure 3A). Of note, downregulation of WRNIP1 significantly reduced the strong ATM activation in WS cells (Figure 3A), confirming that WRNIP1 is required in establishing the idiosyncratic ATM signalling associated with loss of WRN (Marabitti *et al*, 2019).

**Figure 3.**
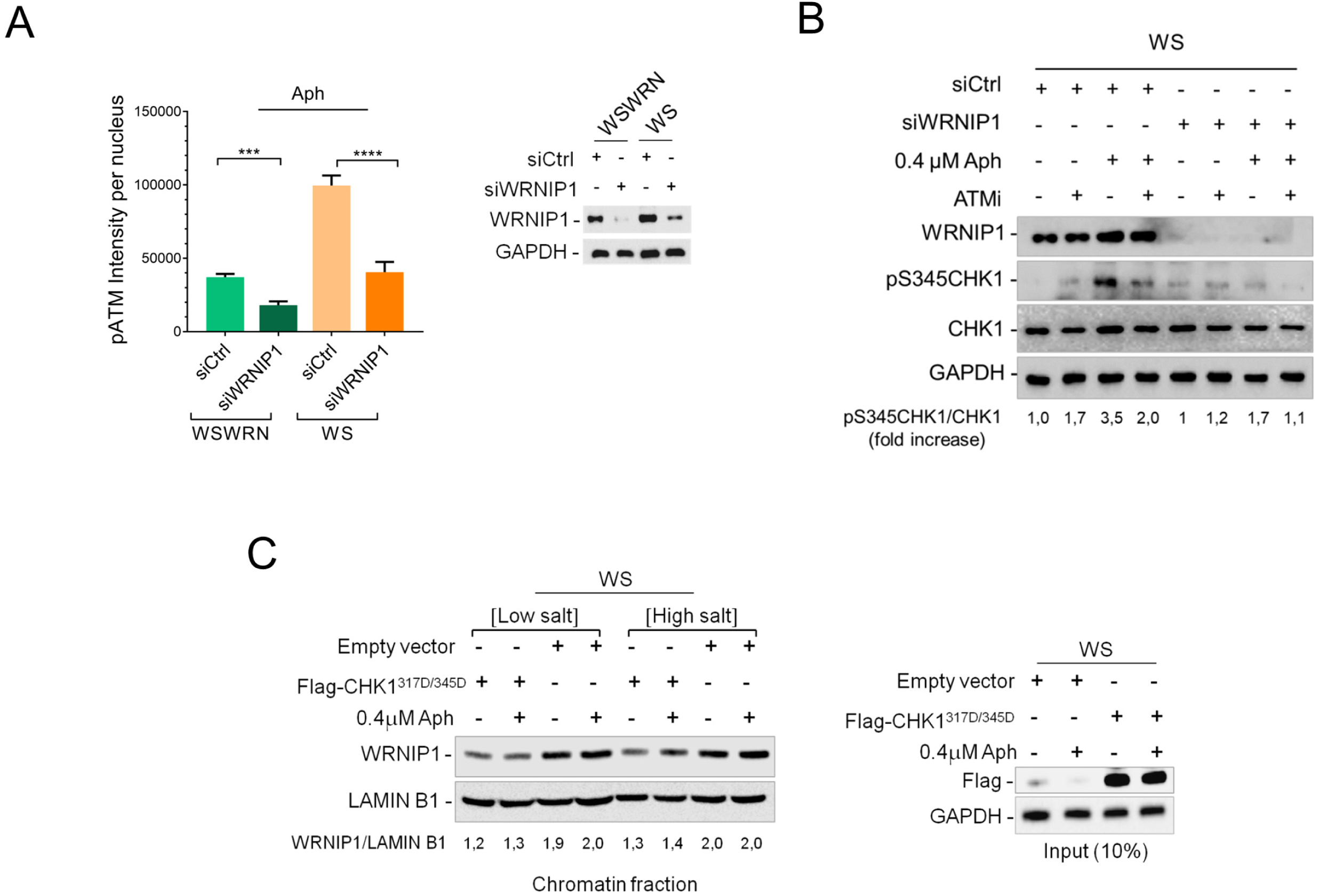
WRNIP1 mediates ATM-dependent CHK1 phosphorylation in WRN-deficient cells. **(A)** Evaluation of ATM activation by immunofluorescence analysis in WSWRN or WS cells depleted for WRNIP1. Cells were transfected with siRNAs directed against GFP (siCtrl) or WRNIP1 (siWRNIP1). After transfection, cells were treated or not with Aph, and subjected to immunostaining using an anti-pATM (S1981) antibody. Bar graph shows pATM intensity per nucleus. Depletion of WRNIP1 was verified by immunoblotting using an anti-WRNIP1 antibody. GAPDH was used as a loading control. Error bars represent standard error (*** P < 0.001; **** P < 0.0001; two-tailed Student’s t test). **B)** WB detection of CHK1 activation in total extracts of WS cells depleted for WRNIP1 as in C. After transfection, cells were exposed to ATMi (KU55933) 1 h prior to be treated or not with Aph. The presence of activated, i.e. phosphorylated, CHK1 was assessed using S345 phospho-specific antibody (pS345). Total amount of CHK1 was determined with an anti-CHK1 antibody. WRNIP1 depletion was confirmed using an anti-WRNIP1 antibody. Equal loading was confirmed probing the membrane with an anti-GAPDH antibody. The normalized ratio of the phosphorylated CHK1/total CHK1 is given. **C)** Analysis of chromatin binding of WRNIP1 in WS cells transfected with Flag-empty vector or Flag-tagged CHK1^317/345D^, and treated or not with Aph. Chromatin fractionation was performed as in Fig.2A. The membrane was probed with the anti-WRNIP1 antibody. LAMIN B1 was used as a loading for the chromatin fraction. Expression levels of Flag-CHK1^317/345D^ (Input) were determined by immunoblotting with anti-Flag antibody. Anti-GAPDH antibody was used to assess equal loading. The normalized ratio of the WRNIP1/LAMIN B1 signal (chromatin) is reported.

Next, we tested whether the WRNIP1-mediated ATM pathway could participate to late CHK1 activation observed in WS cells (Marabitti *et al*, 2019). To this aim, WS cells were depleted for WRNIP1 and treated or not with Aph. Loss of WRNIP1 compromised CHK1 activation after treatment in WS cells, and the reduction of phospho-CHK1 levels was similar to that caused by ATM inhibition (Figure 3B). Consistently with a role of WRNIP1 in activating an ATM signalling, combined loss of WRNIP1 and ATM did not have any additive effect on CHK1 phosphorylation (Figure 3B). Absence of WRN differently affects CHK1 activation upon mild replication stress: it reduces the ATR-dependent CHK1 activation early after Aph treatment but stimulates that dependent on ATM at late time-points (Marabitti *et al*, 2019). The two phenotypes are interlinked and expression of a phospho-mimic form of CHK1, CHK1^317D/345D^ (Gatei *et al*, 2003), prevents the late phenotype (Marabitti *et al*, 2019). Hence, we verified whether expression of the phospho-mimic mutant of CHK1 could hinder stable association of WRNIP1 with chromatin in WS cells. Notably, a normal binding of WRNIP1 to chromatin was restored by expression of the phospho-mimic CHK1 mutant in WS cells (Figure 3C). This suggests that WRNIP1 is required for an ATM-mediated activation of CHK1 and that its stable recruitment in chromatin is triggered by the impaired early CHK1 phosphorylation.

To reinforce this hypothesis, we investigated the binding of WRNIP1 to chromatin in WRN^K577M^ cells, which efficiently phosphorylate CHK1 early after Aph (Basile *et al*, 2014). Fractionation analysis showed that the level of chromatin-bound WRNIP1 in WRN^K577M^ cells was lower than that in WS cells (Suppl. Figure 2A). However, following CHK1 inhibition by UCN-01 (Hui & Helen, 2001), the amount of chromatin-associated WRNIP1 was greatly enhanced in WRN^K577M^ cells as well as in wild-type cells (Suppl. Figure 2B and C).

Therefore, our findings suggest that, in WRN-deficient cells, WRNIP1 is strongly associated with chromatin and related to ATM-dependent CHK1 phosphorylation in response to mild replication stress.

### Depletion of essential factors for activation of ATR-CHK1 pathway promotes WRNIP1 retention in chromatin

Increased stability of WRNIP1 in chromatin is linked to defective CHK1 activation observed in the absence of WRN early after mild replication stress. Several factors facilitate the early ATR-mediated CHK1 phosphorylation, including the ATR kinase-activating protein TopBP1 (Kumagai *et al*, 2006) and the CHK1-interacting factor Claspin (Chini & Chen, 2003; Kumagai *et al*, 2004). Hence, we asked whether increased binding of WRNIP1 to chromatin is a general response to compromised phosphorylation of CHK1. To this aim, we used siRNAs to deplete endogenous TopBP1 or Claspin expression in wild-type cells (WSWRN) and examined WRNIP1 retention in chromatin at high-salt concentration upon mild replication stress. The total amount of WRNIP1 was comparable in mock-depleted, TopBP1- or Claspin-depleted cells under unperturbed conditions or after treatment (Figure 4A). Interestingly, however, TopBP1 or Claspin depletion greatly enhanced WRNIP1 binding to chromatin under high-salt concentration irrespective of the Aph treatment (Figure 4A).

**Figure 4.**
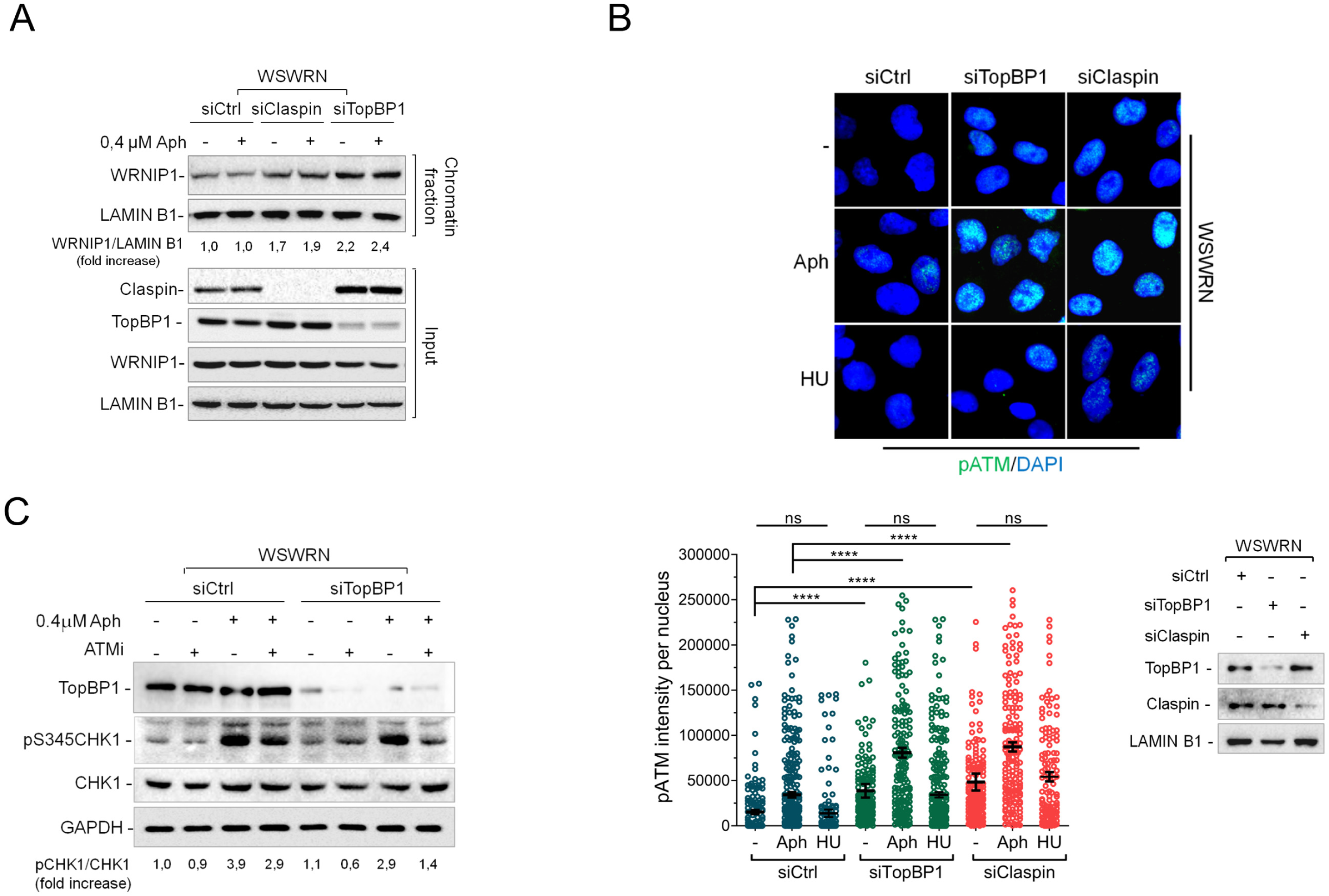
Impairment of ATR-dependent checkpoint triggers WRNIP1-dependent ATM activation. **A)** Analysis of chromatin binding of WRNIP1 in WSWRN cells transfected with siRNAs directed against GFP (siCtrl) or Claspin (siClaspin) or TopBP1(siTopBP1), and treated or not with Aph. Chromatin fractionation was performed under high salts concentration as reported in the experimental scheme in Fig.2. The membrane was probed with an anti-WRNIP1 antibody. LAMIN B1 was used as a loading for the chromatin fraction. Depletion of Claspin or TopBP1was verified by immunoblotting using the relevant antibodies. Total amount of WRNIP1 (Input) in the cells was determined with the relevant antibody. GAPDH was used as a loading control. The normalized ratio of the chromatin-bound WRNIP1/LAMIN B1 is reported. **B)** Evaluation of ATM activation by immunofluorescence analysis in WSWRN cells depleted of Claspin or TopBP1 as in (A). After transfection, cells were treated or not with Aph, and subjected to immunostaining using an anti-pATM (S1981) antibody. Scatter dot plot shows pATM intensity per nucleus. Error bars represent standard error (ns, not significant; **** P < 0.0001; two-tailed Student’s t test). Depletion of Claspin or TopBP1 was verified by immunoblotting using the relevant antibodies. GAPDH was used as a loading control. **C)** WB detection of CHK1 activation in total extracts of WSWRN cells depleted of TopBP1 as in (A). After transfection, cells were exposed to ATMi (KU55933) 1 h prior to be treated or not with Aph. The presence of activated, i.e. phosphorylated, CHK1 was assessed using S345 phospho-specific antibody (pS345). Total amount of CHK1 was determined with an anti-CHK1 antibody. TopBP1 depletion was confirmed using an anti-TopBP1 antibody. Equal loading was confirmed probing the membrane with an anti-GAPDH antibody. The normalized ratio of the phosphorylated CHK1/total CHK1 is given.

Loss of WRN leads to a significant induction of phospho-ATM upon mild replication stress in wild-type cells (Marabitti *et al*, 2019). We therefore evaluated the phosphorylation of ATM in wild-type cells depleted of TopBP1 or Claspin. Staining against pATM showed that, after depletion of TopBP1 or Claspin, ATM was hyper-phosphorylated and that Aph significantly increased its activation (Figure 4B). Interestingly, no significant differences were noted using hydroxyurea (HU) as replication-perturbing treatment (Figure 4B), suggesting that this response is specific of a mild replication stress that does not arrest completely replication fork progression.

Next, we explored whether the ATM pathway is involved in activating CHK1 in TopBP1-depleted cells as observed in WS cells. Our results showed that, although Aph activated CHK1 in both cell lines, CHK1 phosphorylation was only modestly reduced by ATM inhibition in wild-type cells but appeared considerable hampered in TopBP1-depleted cells (Figure 4C). As expected, CHK1 was not phosphorylated after short term exposure to Aph in the absence of TopBP1 (Suppl. Figure 3).

Collectively, these results indicate that impairment of the early ATR-dependent CHK1 activation after mild replication stress calls for increased stability of WRNIP1 in chromatin. Furthermore, they suggest that a WRNIP1-mediated ATM signalling is hyper-activated whenever CHK1 phosphorylation is defective.

### Combined abrogation of ATM activity and WRNIP1 enhances DNA damage after mild replication stress in cells with defective ATR-CHK1 signalling

Our data indicate that mild replication stress triggers a WRNIP1-mediated ATM signalling in cells with impaired CHK1 phosphorylation. However, beside its role in promoting an ATM pathway, WRNIP1 acts as a replication fork protection factor (Leuzzi *et al*, 2016). Moreover, ATM inhibition exacerbates the already elevated genomic instability in WRN-deficient cells (Marabitti *et al*, 2019). Hence, we asked whether abrogating the ATM pathway and WRNIP1 function could exacerbate genome instability further in cells with defective ATR-CHK1 signalling in response to mild replication stress. As model of impaired early activation of CHK1 we used WS cells or cells depleted of TopBP1 or Claspin. We first performed the alkaline Comet assay in WS cells treated with ATM inhibitor and/or siRNA against WRNIP1. As expected, in the absence of WRN, depletion of ATM or WRNIP1 markedly increased the extent of DNA damage after Aph and concomitant depletion of ATM/WRNIP1 further strengthened it (Figure 5A).

**Figure 5.**
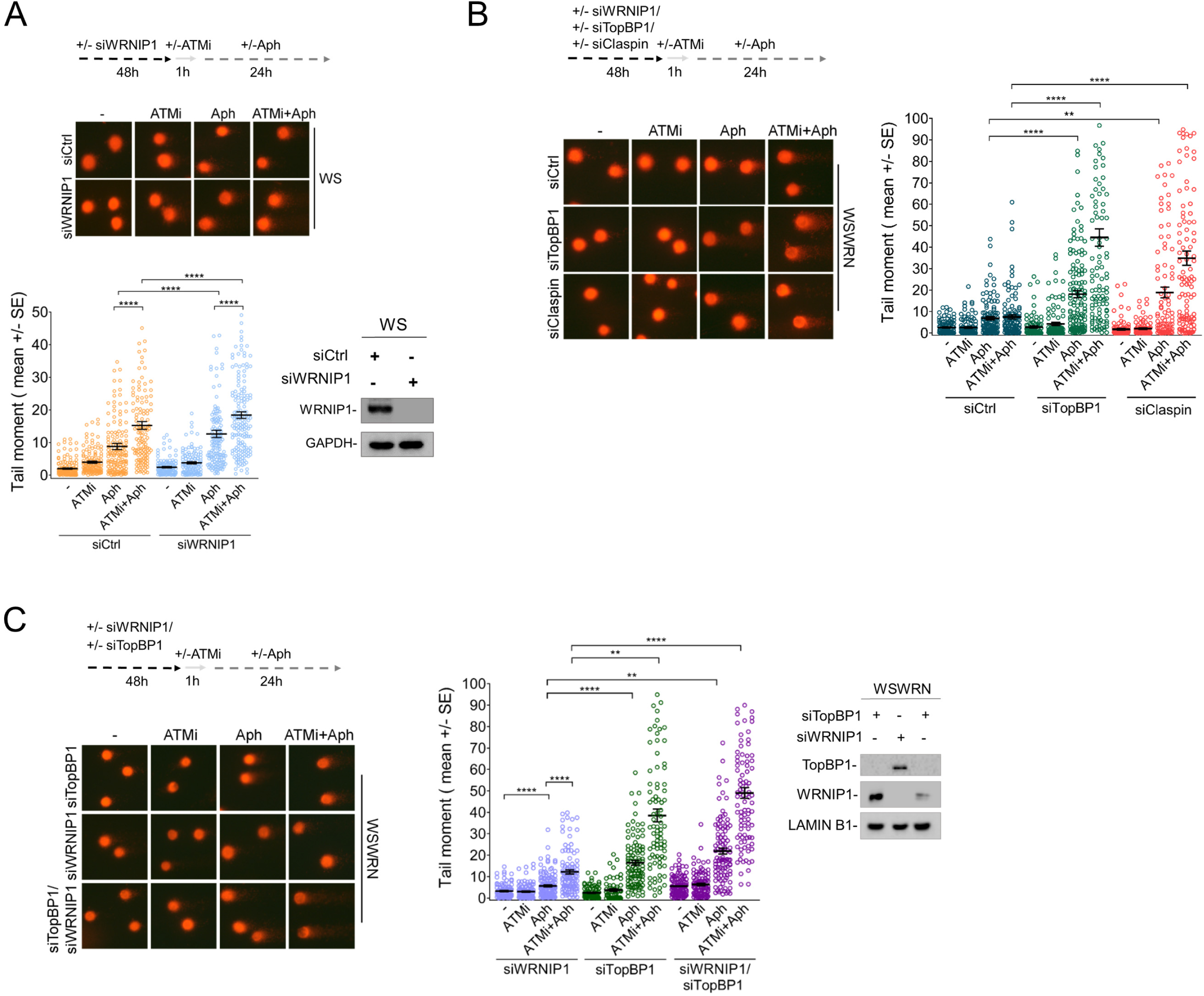
Defects in the ATR-dependent checkpoint sensitize cells to combined abrogation of ATM activity and WRNIP1. **A)** Analysis of DNA damage accumulation evaluated by alkaline Comet assay in WS cells transfected with control siRNAs (siCTRL) or siRNA against WRNIP1, and subjected to chemical inhibition of ATM and/or Aph treatment, as reported in the experimental scheme. Representative images are given. Graph shows data presented as tail moment +/- SE from three independent experiments. Horizontal black lines represent the mean (**** P < 0.0001; two-tailed Student’s t test). **B)** Evaluation of DNA breakage accumulation after TopBP1 or Claspin depletion in WSWRN cells by alkaline Comet assay as reported in the experimental scheme. Representative images are given. Graph shows data presented as tail moment +/- SE from three independent experiments. Horizontal black lines represent the mean (** P < 0.01; **** P < 0.0001; two-tailed Student’s t test). **C)** Analysis of the effect of TopBP1 and/or WRNIP1 depletion on DNA damage accumulation evaluated by alkaline Comet assay in WSWRN as reported in the experimental scheme. Representative images are given. Graph shows data presented as tail moment +/- SE from three independent experiments. Horizontal black lines represent the mean (** P < 0.01; **** P < 0.0001; two-tailed Student’s t test).

Similarly, the downregulation of TopBP1 or Claspin in WSWRN cells resulted in DNA damage potentiation upon mild replication stress (Figure 5B). Notably, more than an additive effect was detected after a combination of ATM inhibition and TopBP1 or Claspin depletion in Aph-treated wild-type cells (Figure 5B).

Of note, treatment with Aph induced a significantly higher level of DNA breakage in cells with concomitant loss of ATM/WRNIP1 than in cells depleted of WRNIP1 alone (Figure 5C). However, upon mild replication stress, the amount of DNA damage was even further increased in wild-type cells with triple TopBP1/ATM/WRNIP1 abrogation in comparison with double knockdown (Figure 5C).

Therefore, combined loss of ATM activity and WRNIP1 potentiates DNA damage after mild replication stress in cells with defective ATR-CHK1 signalling. This suggests that WRNIP1 may play an additional role beyond its function in activating an ATM pathway.

### Retention of WRNIP1 in chromatin correlates with the presence of RAD51 in WS cells

We have previously demonstrated that WRNIP1 stabilises RAD51 on HU-induced replication arrest (Leuzzi *et al*, 2016). We therefore investigated whether WRNIP1 and RAD51 is correlated upon mild replication stress. To this end, we performed a chromatin fractionation assay in wild-type (WSWRN) and WRN-deficient (WS) cells subjected to low-dose Aph at early or late time-points. As expected, under unperturbed conditions and after 24 hours of treatment, high-salt concentration weakened the binding of WRNIP1 to chromatin in wild-type but not in WS cells (Figure 6A). Similar results were obtained at 8 hours of Aph (Figure 6A). Importantly, in WRN-deficient cells, the high levels of WRNIP1 correlated with the presence of elevated amounts of RAD51 (Figure 6A). In agreement with this observation, depletion of WRNIP1 abolished RAD51 retention in chromatin in WS cells (Figure 6B). Further supporting our hypothesis, inhibition of CHK1 activity led to an accumulation of WRNIP1 and consequently of RAD51 in wild-type cells (Figure 6C and Suppl. Figure 2C). By contrast, overexpression of a phospho-mimic mutant of CHK1 promoted removal of both proteins from chromatin in WS cells (Figure 6D).

**Figure 6.**
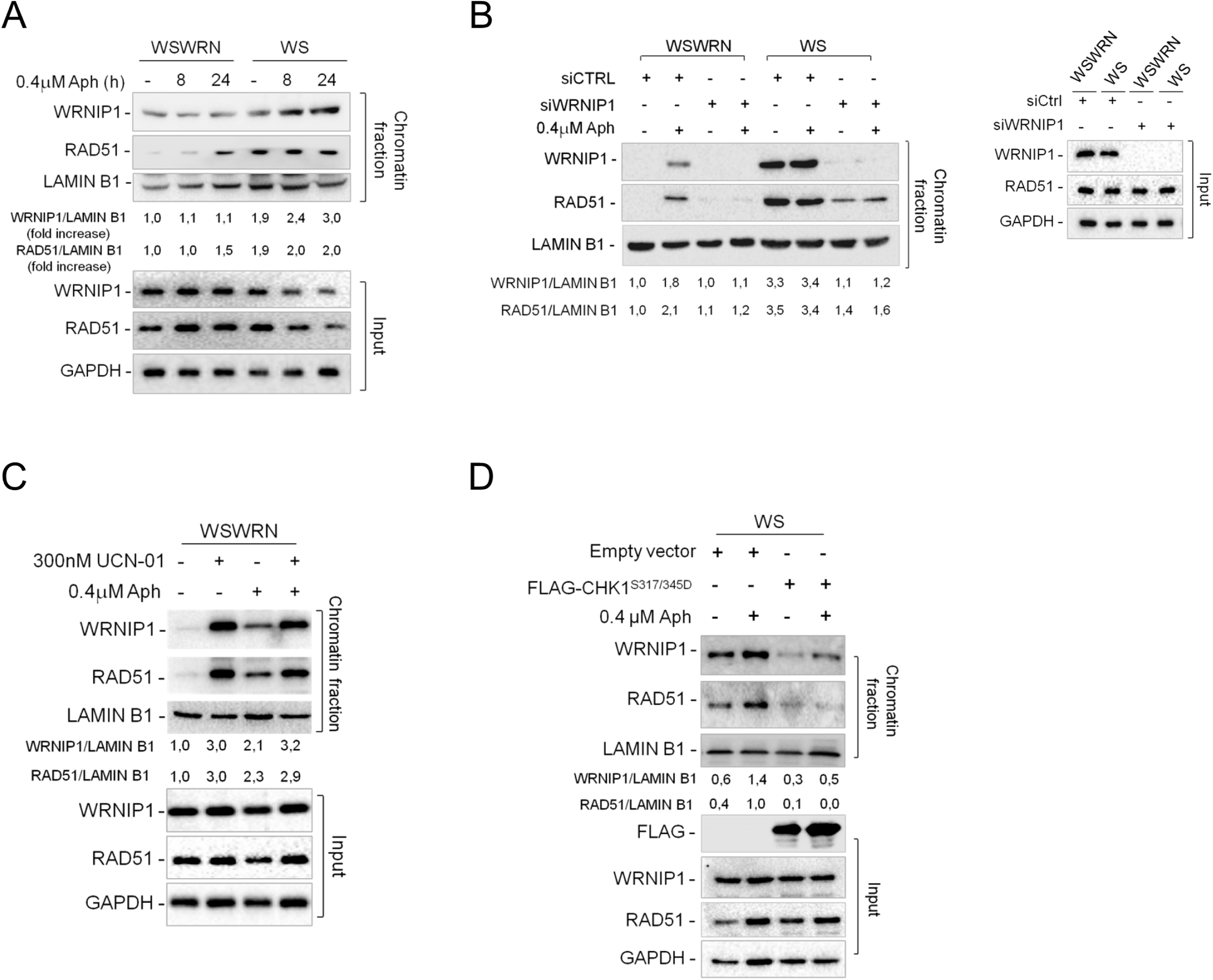
WRNIP1 stabilises RAD51 onto chromatin fraction in WS cells. **A)** Analysis of chromatin binding of WRNIP1 and RAD51 in WSWRN and WS cells treated or not with Aph at 8 and 24 h time-points. Chromatin fractionation was performed under high salts concentration as reported in the experimental scheme in Fig.2A. The membrane was probed with anti-WRNIP1 and anti-RAD51 antibodies. LAMIN B1 was used as a loading for the chromatin fraction. Total amount of WRNIP1 and RAD51 (Input) in the cells was determined with the relevant antibodies. GAPDH was used as a loading control. The normalized ratio of the chromatin-bound WRNIP1 or RAD51/LAMIN B1 is reported. **B)** Analysis of chromatin binding of RAD51 in WSWRN and WS transfected with siRNAs directed against GFP (siCtrl) or against WRNIP1 (siWRNIP1). Cells were treated or not with Aph for 24 hours, then processed for chromatin fractionation as above. The membrane was probed with anti-RAD51 antibody. LAMIN B1 was used as a loading for the chromatin fraction. Total amount of WRNIP1 and RAD51 (Input) in the cells was determined with the relevant antibodies. GAPDH was used as a loading control. The normalized ratio of the chromatin-bound WRNIP1 or RAD51/LAMIN B1 is reported. **C)** Evaluation of the effect of CHK1 inhibition on WRNIP1 and RAD51 recruitment onto chromatin in WSWRN cells. WSWRN cells were exposed or not to Aph, and/or to 300 nM of CHK1 inhibitor, UCN-01, for the last 6 h. Chromatin fractionation of cell lysates was performed as above. The membrane was probed with anti-WRNIP1 and anti-RAD51 antibodies. LAMIN B1 was used as a loading for the chromatin fraction. Total amount of WRNIP1 and RAD51 (Input) in the cells was determined with the relevant antibody. GAPDH was used as a loading control. The ratio of the chromatin-bound WRNIP1 normalized to the untreated is reported below each line. **D)** Analysis of chromatin binding of WRNIP1 and RAD51 in WS cells transfected with Flag-tagged CHK1^317/345D^, and treated or not with Aph. Chromatin fractionation was performed under high salts concentration as in Fig.2A. The membrane was probed with anti-WRNIP1 and anti-RAD51 antibodies. LAMIN B1 was used as a loading for the chromatin fraction. Expression levels of FLAG-CHK1^317/345D^ (Input) were determined by immunoblotting with anti-FLAG antibody. Total amount of WRNIP1 and RAD51 (Input) in the cells was determined with the relevant antibodies. GAPDH was used as a loading control. Anti-GAPDH antibody was used to assess equal loading. The ratio of the WRNIP1/LAMIN B1 signal (chromatin) is reported.

Overall, our observations suggest that WRNIP1 may play a role in stabilising RAD51 after mild replication stress in WRN-deficient cells because of the impaired early CHK1 activation.

### WRNIP1 stimulates the association of RAD51 with ssDNA in proximity of R-loops upon mild replication stress in WS cells

WRNIP1 protects stalled replication forks from degradation promoting RAD51 stabilization on ssDNA (Leuzzi *et al*, 2016). Hence, we first verified whether ssDNAs accumulate upon mild replication stress in WS cells. WSWRN and WS cells were pre-labelled with the thymidine analogue 5-iodo-2’-deoxyuridine (IdU) and treated with Aph for different times as described in the scheme (Figure 7A). We specifically visualized ssDNA formation at parental-strand by immunofluorescence using an anti-BrdU/IdU antibody under non-denaturing conditions as reported (Basile *et al*, 2014). Our analysis showed that WRN-deficient cells presented a significant higher amount of ssDNA than wild-type cells at later time-points of treatment (Figure 7A).

**Figure 7.**
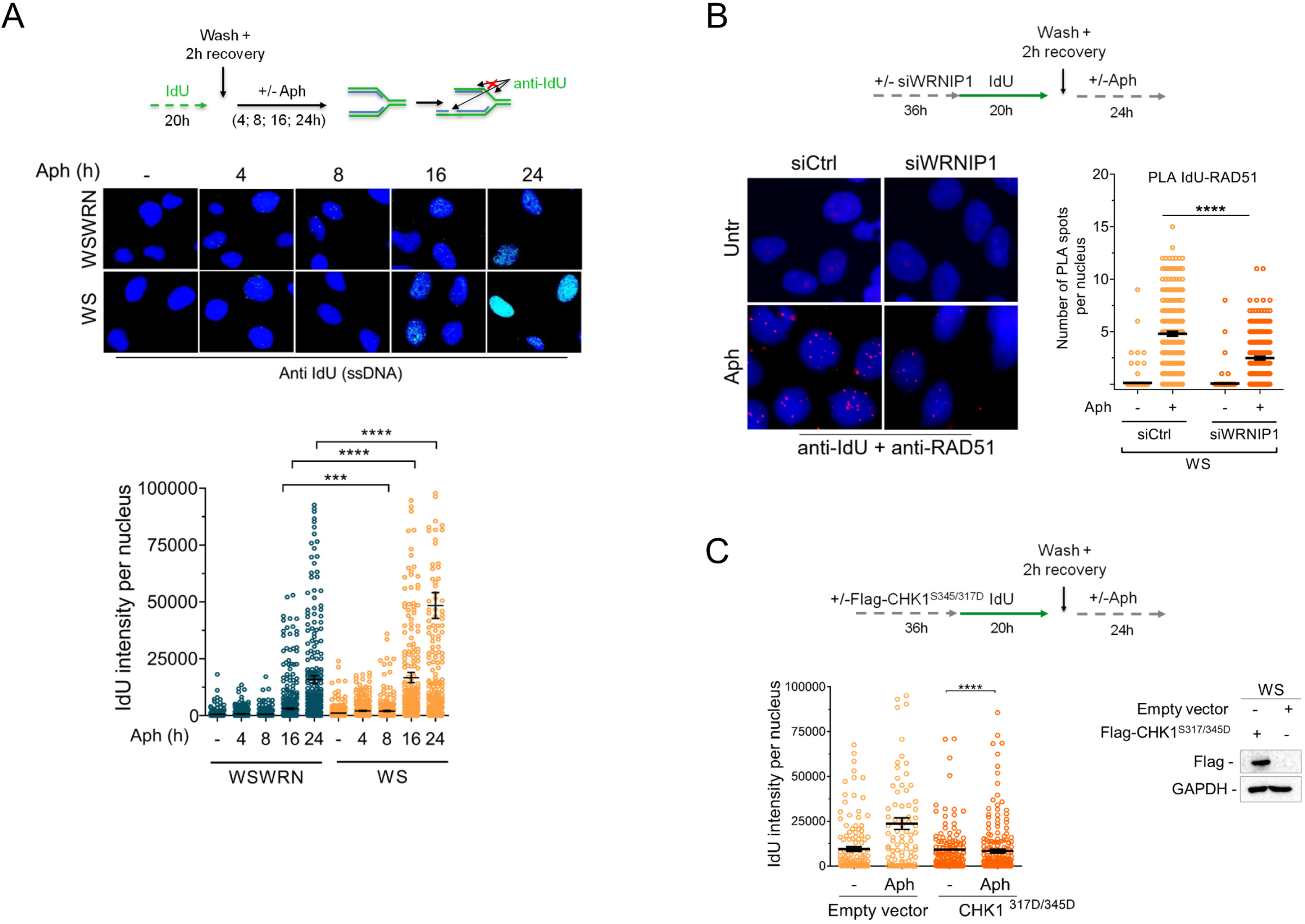
WRNIP1 promotes RAD51 stabilisation onto ssDNA gaps generated by prolonged exposure to Aph in WS cells. **A)** Evaluation of ssDNA accumulation at parental-strand by immunofluorescence analysis in WSWRN or WS cells. Experimental design of ssDNA assay is shown. Cells were labelled with IdU for 24 h, as indicated, washed and left to recover for 2 h, then treated or not with 0.4 µM Aph for 4, 8, 16, 24 h. After treatment, cells were fixed and stained with an anti-IdU antibody without denaturing the DNA to specifically detect parental ssDNA. Horizontal black lines and error bars represent the mean +/- SE; n = 3 (***P < 0.001; ****P < 0.0001; two-tailed Student’s t-test). Representative images are shown. DNA was counterstained with DAPI (blue). **B)** Analysis of DNA-protein interactions between ssDNA and endogenous RAD51 in WS cells depleted of WRNIP1 by *in situ* PLA assay. Experimental designed used for the assay is given. Cells were transfected as reported and labelled with IdU as above. Next, cells were fixed, stained with an anti-IdU antibody and subjected to PLA assay as described in the “Materials and methods” section. Antibodies raised against IdU or RAD51 were used to reveal ssDNA or endogenous RAD51 respectively. Each red spot represents a single interaction between ssDNA and RAD51. No spot has been revealed in cells stained with each single antibody (negative control). DNA was counterstained with DAPI (blue). Representative images of the PLA assay are given. Graph shows the number of PLA spots per nucleus. Horizontal black lines represent the mean value (****, P < 0.0001; two-tailed Student’s t test). **C)** Evaluation of ssDNA accumulation at parental-strand by immunofluorescence analysis in WS cells transfected with an empty vector or a Flag-tagged CHK1^317/345D^, and treated or not with Aph 48 h post-transfection. Experimental design of ssDNA assay is shown. Expression levels of FLAG-CHK1317/345D were determined by immunoblotting with anti-Flag antibody. Anti-GAPDH antibody was used to assess equal loading. Horizontal black lines and error bars represent the mean +/- SE; n = 3 (****P < 0.0001; two-tailed Student’s t-test). Representative images are shown. DNA was counterstained with DAPI (blue).

Next, we wondered if RAD51 localises on parental ssDNA in a WRNIP1-dependent manner. Using a modification of the in situ proximity ligation assay (PLA), a fluorescence-based improved method that makes possible to detect protein/DNA association (Leuzzi *et al*, 2016; Malacaria *et al*, 2019; Iannascoli *et al*, 2015), we investigated the co-localisation of RAD51 at/near ssDNA. To do this, WS cells depleted of WRNIP1 by RNAi were treated or not with Aph (Figure 7B). We found that the co-localisation between ssDNA (anti-IdU signal) and RAD51 significantly increased after mild replication stress in WS cells (Figure 7B). By contrast, and in agreement with the reduced levels of RAD51 in chromatin, less PLA spots were observed in the absence of WRNIP1 (Figure 7B). This result suggests that, in WS cells, the RAD51-ssDNA association requires WRNIP1 also after mild replication stress.

It is important to note that, in WS cells, the amount of ssDNA decreased upon mild replication stress by transfecting a phospho-mimic mutant of CHK1, a condition that abolishes the need of the ATM pathway activation or WRNIP1 and RAD51 retention in chromatin (Figure 7C).

As loss of WRN results in R-loop accumulation that is responsible for ATM signalling activation (Marabitti *et al*, 2019), we assessed if ssDNA could arise from persistent DNA-RNA hybrids. To prove this, we analysed the effect of overexpression of ectopic GFP-RNaseH1, a ribonuclease that degrades RNA engaged in R-loops (Nguyen *et al*, 2017), on ssDNA formation. As can be seen in Fig. 8A, the IdU intensity per nucleus was significantly suppressed by RNaseH1 overexpression in both the WSWRN and WS cell lines with or without treatment. Consistently, degradation of R-loops counteracted retention in chromatin of WRNIP1 and RAD51 in WS cells (Figure 8B). By contrast, preventing the processing of R-loops into DSBs by the endonuclease XPG heightened slightly the levels of parental ssDNA in WS cells, arguing against an end-resection-related origin (Suppl. Figure 4).

**Figure 8.**
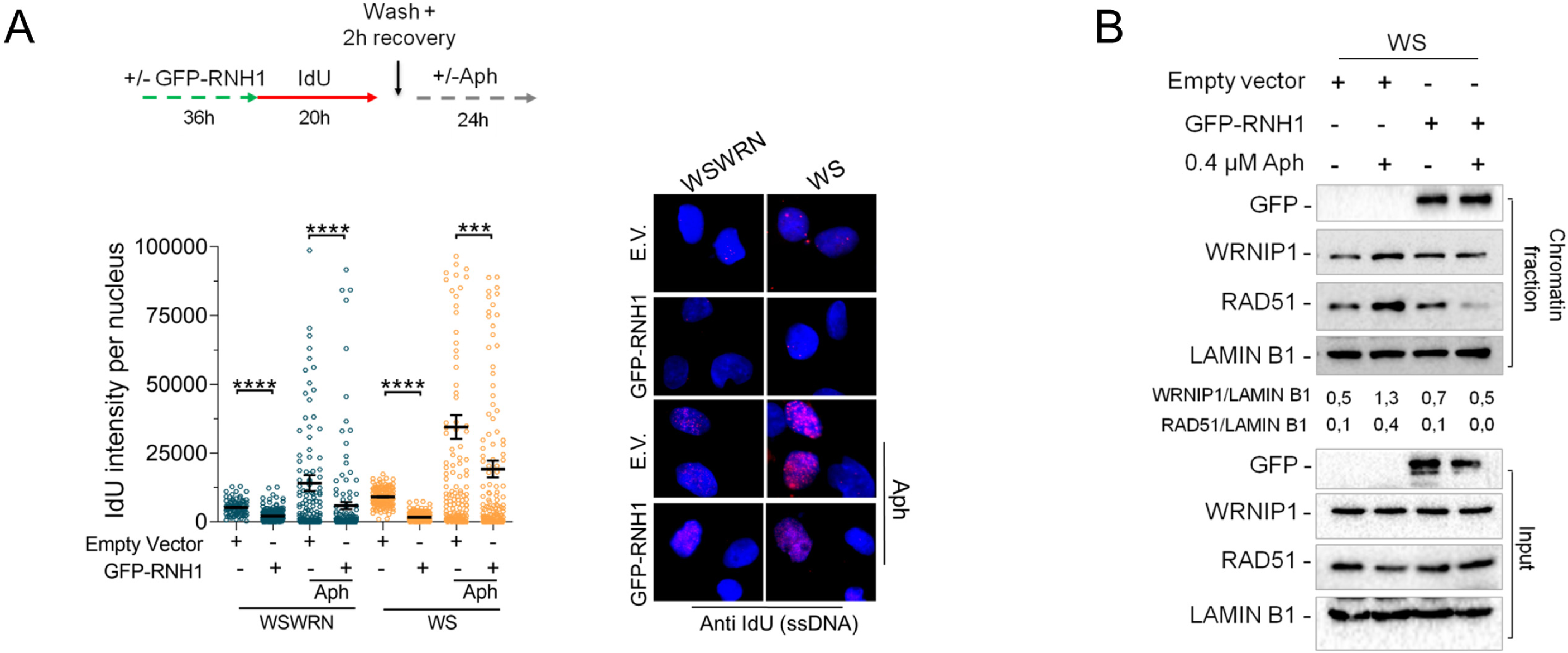
R-loops accumulation leads to ssDNA exposure requiring WRNIP1 protective function in WRN-deficient cells. **A)** Evaluation of ssDNA accumulation at parental-strand by immunofluorescence analysis in WSWRN or WS cells treated as reported in the experimental scheme after transfection of a vector expressing GFP-tagged RNaseH1. Horizontal black lines and error bars represent the mean +/- SE; n = 3 (***P < 0.001,****P < 0.0001; two-tailed Student’s t-test). Representative images are shown. DNA was counterstained with DAPI (blue). **B)** Evaluation of the effect of RNAseH1 over-expression on WRNIP1 and RAD51 recruitment on chromatin. WS cells were transfected as in A) and treated or not with Aph 48h post-transfection. Chromatin fractionation was performed under high salts concentration as described in Fig.2A. The membrane was probed with anti-WRNIP1 and anti-RAD51 antibodies. LAMIN B1 was used as a loading for the chromatin fraction. Expression levels of GFP-RNH1 (Input) were determined by immunoblotting with anti-GFP antibody. Total amount of WRNIP1 and RAD51 (Input) in the cells was determined with the relevant antibodies. GAPDH was used as a loading control. Anti-GAPDH antibody was used to assess equal loading. The ratio of the WRNIP1/LAMIN B1 signal (chromatin) is reported.

To extend the above observations to the cells with defective ATR-CHK1 signalling, we evaluated direct accumulation of R-loops by immunofluorescence in WSWRN cells in which TopBP1 or Claspin was depleted. Our analysis showed that spontaneous levels of S9.6 nuclear intensity in both TopBP1- and Claspin-depleted cells were significantly higher than those in wild-type cells (Figure 9A). A significant enrichment of the S9.6 nuclear signal was observed upon Aph-treatment in cells depleted for TopBP1 or Claspin respect to wild-type counterpart (Figure 9A).

**Figure 9.**
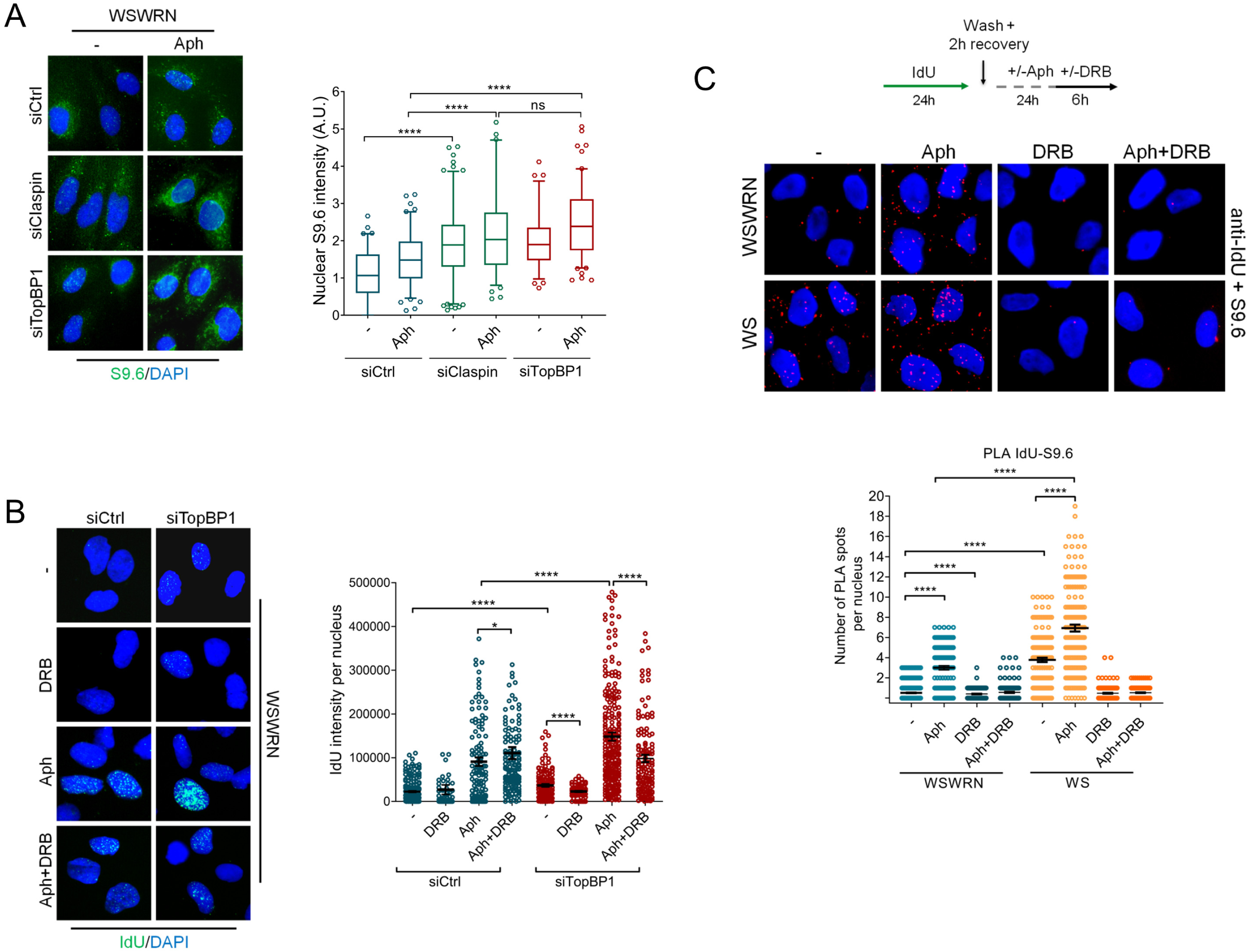
Depletion of checkpoint mediators cause R-loop accumulation leading to ssDNA gaps formation. **A)** Evaluation of R-loops accumulation by immunofluorescence analysis. WSWRN cells transfected with siRNAs directed against GFP (siCtrl) or Claspin (siClaspin) or TopBP1(siTopBP1), and treated or not with Aph. Next, cells were fixed and stained with anti-RNA-DNA hybrids S9.6 monoclonal antibody. Nuclei were counterstained with DAPI. Representative images are given. Box plot shows nuclear S9.6 fluorescence intensity. Box and whiskers represent 20-75 and 10-90 percentiles, respectively. Horizontal line represents the mean value. Error bars represent standard error (**** P < 0.0001; two-tailed Student’s t test). **B)** Evaluation of transcription inhibition on ssDNA accumulation at parental-strand by immunofluorescence analysis in WSWRN cells transfected with control siRNAs (siCtrl) or siRNAs directed against Topbp1 (siTopBP1). After 48 h cells were treated as reported in the experimental scheme. After treatment, cells were fixed and stained with an anti-IdU antibody without denaturing the DNA to specifically detect parental ssDNA. Horizontal black lines and error bars represent the mean +/- SE; n = 3 (* P < 0.1; **** P < 0.0001 two-tailed Student’s t-test). Representative images are shown. DNA was counterstained with DAPI (blue). **C)** Analysis of interactions between ssDNA and R-loops in WSWRN and WS cells by *in situ* PLA assay. Experimental designed used for the assay is given. Cells were labelled with IdU as above and treated or not with Aph and/or DRB. Next, cells were fixed, stained with an anti-IdU antibody and subjected to PLA assay as described in the “Materials and methods” section. Antibodies raised against IdU or R-loops (S9.6) were used to reveal parental ssDNA or nuclear R-loops respectively. Each red spot represents a single interaction between ssDNA and R-loops. No spot has been revealed in cells stained with each single antibody (negative control). DNA was counterstained with DAPI (blue). Representative images of the PLA assay are given. Graph shows the number of PLA spots per nucleus. Horizontal black lines represent the mean value (****, P < 0.0001; two-tailed Student’s t test).

In addition, TopBP1-depleted cells accumulated ssDNA upon mild replication stress, but treatment with 5, 6-dichloro-1-ß-D-ribofurosylbenzimidazole (DRB), an inhibitor of RNA elongation (Salas-Armenteros *et al*, 2017), led to a strong reduction of the IdU intensity per nucleus (Figure 9B).

Finally, to further confirm that ssDNAs are formed at/near R-loops, we performed a PLA assay. WSWRN and WS cells were incubated with Aph and DRB, then subjected to PLA using an anti-BrdU/ldU (ssDNA) and an anti-DNA-RNA hybrids S9.6 (R-loop) antibodies. Our analysis showed an increased number of spontaneous PLA spots in WS cells that were abolished by transcription inhibition (Figure 9C). After Aph treatment, PLA spots were significantly enhanced in both cell lines, with values more elevated in WRN-deficient cells, and almost completely suppressed by DRB (Figure 9C). These evidences indicates a spatial proximity between ssDNAs and R-loops.

Altogether, our findings suggest that RAD51-ssDNA association and accumulation is R-loop-dependent in WRN-deficient cells.

## DISCUSSION

Accumulation of unscheduled R-loops represents a common source of replication stress and genome instability (García-Muse & Aguilera, 2019; Allison & Wang, 2019). Given the negative impact of aberrant R-loops on transcription, replication and DNA repair, cells possess several mechanisms to prevent or resolve such harmful intermediates (García-Muse & Aguilera, 2019; Crossley *et al*, 2019). Currently, it is thought that an important role in avoiding deleterious consequences of these R-loops is played by some DDR proteins and replication fork protection factors (Bhatia *et al*, 2014). Furthermore, it is recently emerged that loss of WRN, a protein involved in the repair and recovery of stalled replication forks, leads to an ATM-pathway activation to limit R-loop-associated genome instability in human cells (Marabitti *et al*, 2019). In this study, we demonstrate that the WRN-interacting protein 1 (WRNIP1) is implicated in the response to R-loop-induced DNA damage in cells with dysfunctional replication checkpoint.

WRNIP1 is a member of the AAA+ ATPase family that was first identified as an interactor of WRN (Kawabe Yi *et al*, 2001), but there is no evidence of a functional relationship between these proteins in response to mild replication stress. Although our results show that WRNIP1 co-immunoprecipitates with WRN under unperturbed conditions and that low-dose of aphidicolin slightly enhances this interaction, we notice that WRNIP1 is essential in the absence of WRN to counteract the effects of unscheduled R-loops. Indeed, loss of WRN or WRNIP1-depletion sensitises human cells to Aph-treatment, but concomitant lack of WRNIP1 and WRN results in a synergistic enhancement of the chromosomal aberrations frequency and DNA damage levels. These observations agree with those obtained from chicken DT40 cells that confirmed the binding of WRNIP1 to WRN, but concomitantly showed that the two proteins function independently to deal with DNA lesions during replication (Kawabe *et al*, 2006). Of note, in yeast, deletion of Mgs1, the homolog of WRNIP1, leads to growth defects and elevated genomic instability, and exhibits a relation of synthetic lethality with Sgs1, the unique bacterial RecQ helicase (Branzei *et al*, 2002).

WRNIP1 is stabilised in chromatin in WS cells and stabilisation correlates with inability to properly activate CHK1 upon aphidicolin, which is consistent with loss of WRN affecting ATR checkpoint activation upon mild replication stress (Marabitti *et al*, 2019; Basile *et al*, 2014). However, WRNIP1 is stabilised in chromatin also upon depletion of TopBP1, which is a key mediator of the ATR kinase (Kumagai *et al*, 2006), indicating that whenever the ATR-CHK1 signalling is dysfunctional WRNIP1 is hyper-activated and retained stably in chromatin. In WS cells, inability to activate CHK1 early after Aph correlates to increased R-loop formation (Marabitti *et al*, 2019). ATR-CHK1 pathway has been previously involved in safeguarding genome integrity against aberrant R-loops (Matos *et al*, 2019; Hamperl *et al*, 2017). This agrees with the ability of deregulated R-loops to hamper replication fork progression (Wellinger *et al*, 2006; Gan *et al*, 2011; Tuduri *et al*, 2009), and also with the recent observation that depletion of ATR or CHK1 leads to R-loop-dependent replication fork stalling (Matos *et al*, 2019). Consistently, and in line with other reports (Matos *et al*, 2019; Barroso *et al*, 2019), we see that abrogation of essential factors for the ATR-dependent checkpoint results in high levels of R-loops. It has been previously shown that WRNIP1 is implicated in the efficient activation of ATM in response to stimuli that do not produce DNA breakage (Pichierri *et al*, 2001; Schmidt *et al*, 2014; Kanu *et al*, 2016), and a DSB-independent but R-loop-dependent ATM pathway has been described in quiescent cells (Tresini *et al*, 2015). Indeed, WRN-deficient cells trigger an ATM signalling specifically after Aph-induced replication stress, which is R-loop dependent (Marabitti *et al*, 2019). In keeping with this, we observe an R-loop-dependent hyper-phosphorylation of ATM in all the conditions tested in which the ATR checkpoint was inhibited. Also, we find that chromatin-bound WRNIP1 is related to the late ATM-dependent CHK1 phosphorylation. Supporting this, WRNIP1 recruitment is counteracted by overexpression of a constitutively active CHK1 that, compensating for defective ATR pathway, abolishes the need to activate ATM as well as in WRN helicase dead cells that efficiently phosphorylate CHK1 (Basile *et al*, 2014; Marabitti *et al*, 2019). Interestingly, degradation of R-loops weakens the association of WRNIP1 with chromatin. Hence, it is not surprisingly that, similarly, every time the replication checkpoint is compromised, WRNIP1 retention in chromatin is required for triggering an ATM-CHK1 signalling, which might be engaged in limiting transcription and/or in preventing massive R-loop-associated DNA damage accumulation. Accordingly, ATM inhibition or WRNIP1 abrogation in these pathological contexts is accompanied by increased levels of genomic instability.

Noteworthy, combined loss of ATM activity and WRNIP1 potentiates DNA damage in cells with dysfunctional ATR checkpoint, suggesting additional roles for WRNIP1 beyond its function as a mediator of ATM. We have recently reported that WRNIP1 stabilises RAD51 at perturbed forks after hydroxyurea treatment (Leuzzi *et al*, 2016). Of note, we observe that retention of RAD51 in chromatin in WS cells correlates with the presence of WRNIP1, and both correlate with the accumulation of R-loops. It is known that BRCA2 mediates RAD51 loading on ssDNA (Jensen *et al*, 2010; Moynahan & Jasin, 2010; Liu *et al*, 2010) and is required for R-loop processing (Bhatia *et al*, 2014). WRNIP1 forms a complex with BRCA2/RAD51 (Leuzzi *et al*, 2016). It has been proposed that, BRCA2 together with other proteins, could contribute to prevent collapse and reversal of R-loop-induced stalled forks, avoiding R-loop extension, and promoting fork restart and R-loop dissolution (Bhatia *et al*, 2014). In this regard, it is tempting to speculate that, in cells with replication checkpoint defects, WRNIP1 could act in concert with BRCA2 to stabilise RAD51 on ssDNA generated near/at sites of replication-transcription conflicts.

Interestingly, we observe that defective ATR checkpoint promotes accumulation of parental ssDNA that is dependent on transcription and R-loops. Such parental ssDNA is detected in proximity of R-loops upon mild replication stress. Upon transcription-replication conflicts, exposure of parental ssDNA might derive from the unwound DNA strand at the R-loop or from the fork stalling in front of the R-loop. Hence, WRNIP1 could contribute to stabilise either RAD51 nucleofilaments assembling at the displaced DNA strand of the R-loop or at the parental DNA exposed at the fork. Of note, the fact that, upon replication-transcription collisions, WRNIP1 stimulates the stability of RAD51 at parental strand might suggest that its function is carried out before fork reversal, which is expected to result in formation of RAD51 nucleofilaments at the exposed nascent strand of the reversed arm (Stirling & Hieter, 2017; Taglialatela *et al*, 2017; Malacaria *et al*, 2019). Therefore, the WRNIP1-mediated RAD51 stabilisation in chromatin might be not specific for RAD51 assembled at reversed forks, as described in hydroxyurea-treated cells (Leuzzi *et al*, 2016).

Previous findings reported that the structure-specific nucleases XPF and XPG directly cleave R-loops to promote their resolution (Sollier *et al*, 2014). Mounting evidences suggest that R-loop-induced ATR activation is independent of XPG, but requires the MUS81 endonuclease (Pichierri *et al*, 2001; Matos *et al*, 2019; Chappidi *et al*, 2019). By contrast, in the absence of WRN, XPG-mediated transient DSBs deriving from R-loop processing are responsible for ATM pathway activation (Marabitti *et al*, 2019). This agrees with previous data showing that, in WRN-deficient cells, treatment with low-dose of Aph does not induce MUS81-dependent DSB formation but determines ssDNA accumulation and enhances the number of RAD51 foci (Murfuni *et al*, 2012). Accordingly, WRNIP1 could play two independent functions upon replication-transcription conflicts: stabilisation of RAD51 at R-loops or R-loop-dependent stalled forks, and activation of ATM to repair downstream DSBs derived from the active processing of R-loops and collisions with the forks (Figure 10).

**Figure 10.**
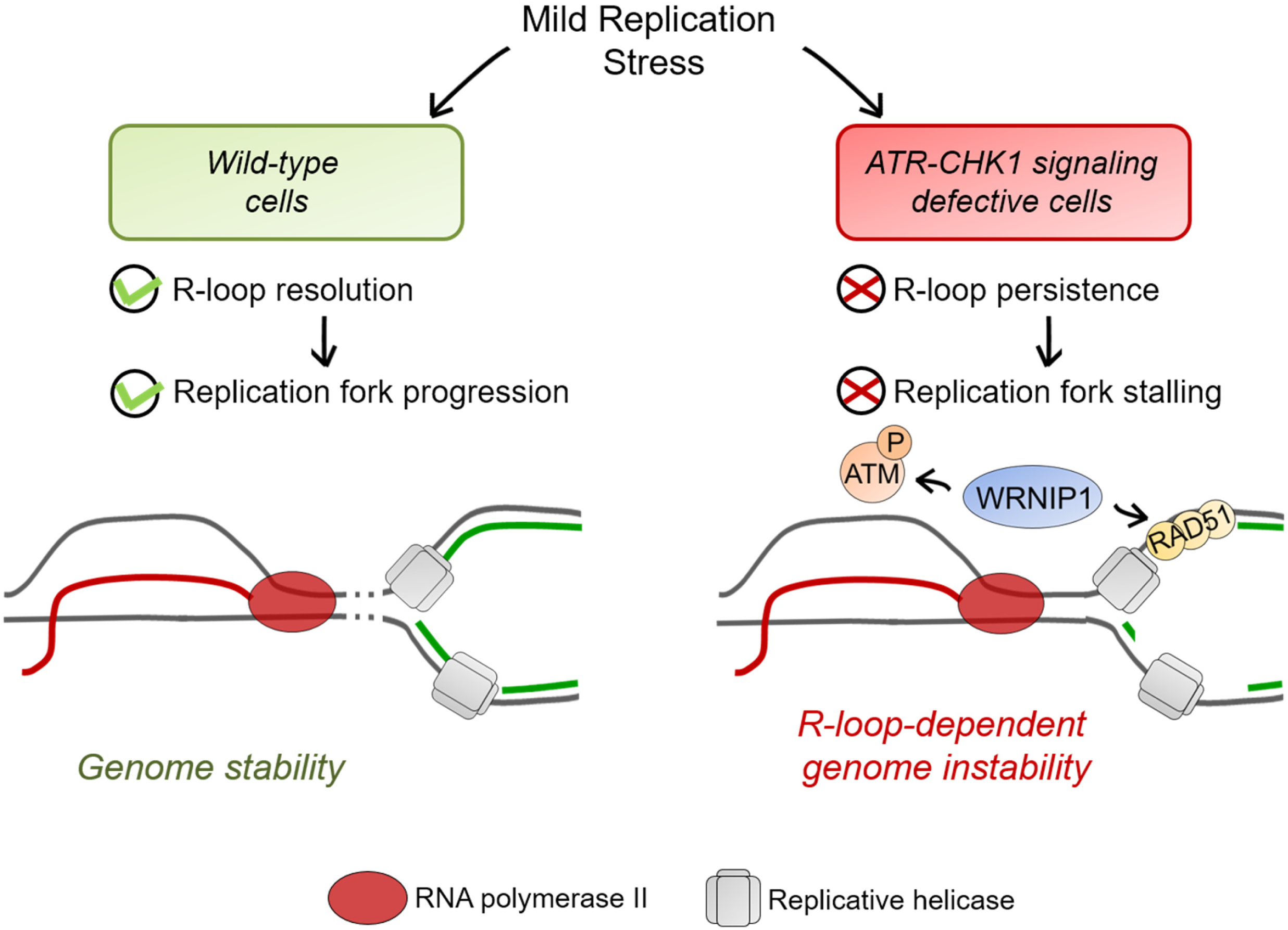
Model for the role of WRNIP1 in the suppression of genome instability derived from transcription-replication conflicts in cells with dysfunctional ATR checkpoint. Under a condition of mild replication stress, cells with proficient ATR checkpoint are able to efficiently prevent R-loop accumulation, thus minimizing transcription-dependent perturbation of DNA replication. By contrast, when ATR-CHK1 signalling is defective, aberrant R-loops accumulate leading to replication fork stalling. Single-stranded DNA gaps generated at transcription-replication conflicts sites require a WRNIP1-mediated response, which triggers an ATM-dependent checkpoint and RAD51-dependent fork protection.

In summary our findings uncover a novel role of the WRNIP1-mediated response in counteracting aberrant R-loop accumulation. Dual function of WRNIP1 is required for proper maintenance of genome stability in the pathological contexts deriving from dysfunctional ATR-dependent checkpoint. As mounting evidences reveal direct connections between R-loops and cancer (Wells *et al*, 2019), the elevated genome instability caused by mild replication stress after WRNIP1 depletion in cells with dysfunctional ATR checkpoint puts forward WRNIP1 as a target to further sensitise cancer cells to inhibitors of ATR or CHK1, which are currently under clinical evaluation.

## AUTHORS CONTRIBUTIONS

V.M. performed fluorescence and biochemical experiments, DNA damage and chromosomal aberrations analyses. G. L. carried out ssDNA experiments. E. M. carried out PLA experiments. V. P. contributed to biochemical experiments. V.M. analysed data and contributed to design experiments. A.F. and P.P designed experiments and analysed data. A.F. and P.P wrote the paper.

## MATERIALS AND METHODS

### Cell cultures

AG11395 (WRN-deficient) human fibroblasts retrovirally-transduced with full length cDNA encoding wild-type WRN (WSWRN) or missense-mutant form of WRN with inactive helicase (WRN^K577M^) were generated as previously described (Pirzio *et al*, 2008). The SV40-transformed MRC5 fibroblast cell line (MRC5SV) was a generous gift from Patricia Kannouche (IGR, Villejuif, France). shWRNIP1 cell line was generated by stably expressing shRNA against WRNIP1 (shWRNIP1) (OriGene). Cells were cultured in the presence of puromycin (100 ng/ml; Invitrogen) to maintain selective pressure for shRNA expression. All cell lines were maintained in DMEM (Invitrogen) supplemented with 10% FBS (Boehringer Manheim) and incubated at 37°C in an humified 5% CO_2_ atmosphere.All the cell lines were maintained in Dulbecco’s modified Eagle’s medium (DMEM; Life Technologies) supplemented with 10% FBS (Boehringer Mannheim), and incubated at 37°C in a humidified 5% CO_2_atmosphere.

### Chromatin fractionation

Chromatin fractionation experiments were performed as previously described with minor modifications (Leuzzi *et al*, 2016). Briefly, 1.5×10^7^ cells were harvested using a cell scraper, centrifuged, and then pellet was washed twice with PBS. Cell pellets were resuspended in buffer A (10 mM HEPES pH 7.9, 10 mM KCl, 1.5 mM MgCl_2_, 0.34 M sucrose, 10% glycerol, 1 mM DTT). Triton X-100 (0.1%) was added, and the cells were incubated for 5 min on ice. Nuclei were collected in pellet by centrifugation. The supernatant was discarded, nuclei washed once in buffer A, and then lysed in buffer B (3 mM EDTA, 0.2 mM EGTA, 1 mM DTT). Insoluble chromatin was collected by centrifugation, washed once in buffer B, and centrifuged again under the same conditions. The chromatin pellet was resuspended in buffer B/CSK (PIPES, 300 mM Sucrose, 3 mM MgCl_2_, 1 mM EGTA) and then split into two equal aliquots. These samples were centrifuged and supernatants discarded. Pellets were resuspended in buffer B/CSK supplemented with 100 mM NaCl ([low salts] extraction), and kept on ice for 10 min. Pellet contains nuclei were subjected to Western blot analysis or to further salt extraction by resuspension in buffer B/CSK supplemented with 300mM NaCl ([high salts] extraction). After an incubation of 10 min on ice, pellets were collected by centrifugation and supernatant was discarded. The final pellets were resuspended in 2× sample loading buffer (100 mM Tris/HCl pH 6.8, 100 mM DTT, 4% SDS, 0.2% bromophenol blue and 20% glycerol), sonicated on ice, boiled for 5 min at 95°C and then subjected to Western blot as reported in “Supplementary Materials and methods”.

### Immunofluorescence

Immunofluorescent detection of phospho-ATM was performed according to standard protocol with minor changes. Briefly, exponential growing cells were seeded onto Petri dishes, then treated (or mock-treated) as indicated, fixed in 3% formaldehyde/2% sucrose for 10 min, and permeabilized using 0.4% Triton X-100 for 10 min prior to incubation with 10% FBS for 1 h. After blocking, cells were incubated with the antibody against phospho-ATM-Ser1981 (Millipore, 1:300) for 2 h at RT. To detect parental-strand ssDNA, cells were pre-labelled for 20 h with 100 µM IdU (SigmaAldrich), washed in drug-free medium for 2 hours and then treated with Aph for 24 h. Next, cells were washed with PBS, permeabilized with 0.5% Triton X-100 for 10 min at 4°C, fixed with 3% formaldehyde/2% sucrose solution for 10 min, and then blocked in 3% BSA/PBS for 15 min as previously described (Couch et al, 2013). Fixed cells were then incubated with anti-IdU antibody (mouse-monoclonal antiBrdU/IdU; clone b44 Becton Dickinson, 1:100). The incubation with antibodies was accomplished in a humidified chamber for 1 h at RT. DNA was counterstained with 0.5 µg/ml DAPI. Images were acquired as described above. Immunostaining for RNA–DNA hybrids was performed as described (Hamperl *et al*, 2017). Briefly, cells were fixed in 100% methanol for 10 min at –20°C, washed three times in PBS, pre-treated with 6 μg/ml of RNase A for 45 min at 37°C in 10 mM Tris–HCl pH 7.5 supplemented with 0.5 M NaCl, before blocking in 2% BSA/PBS overnight at 4°C. Cells were then incubated with the anti-DNA–RNA hybrid [S9.6] antibody (Kerafast) overnight at 4°C. After each primary antibody, cells were washed twice with PBS, and incubated with the specific secondary antibody: goat anti-mouse Alexa Fluor-488 or goat anti-rabbit Alexa Fluor-594 (Molecular Probes). The incubation with secondary antibodies were accomplished in a humidified chamber for 1 h at RT. DNA was counterstained with 0.5 μg/ml DAPI. Images were randomly acquired using Eclipse 80i Nikon Fluorescence Microscope, equipped with a VideoConfocal (ViCo) system. For each time point, at least 200 nuclei were acquired at 40× magnification. Phospho-ATM, IdU or S9.6 intensity per nucleus was calculated using ImageJ. Parallel samples incubated with either the appropriate normal serum or only with the secondary antibody confirmed that the observed fluorescence pattern was not attributable to artefacts.

### Statistical analysis

Statistical differences in all case were determined by two-tailed Student’s *t* test. In all cases, not significant; P > 0.05; *P < 0.05; **P < 0.01; ***P < 0.001; ****P < 0.0001.

